# Parasite genetic variation and systemic immune responses are not associated with different clinical presentations of cutaneous leishmaniasis caused by *Leishmania aethiopica*

**DOI:** 10.1101/2024.04.19.590259

**Authors:** Endalew Yizengaw, Yegnasew Takele, Susanne Franssen, Bizuayehu Gashaw, Mulat Yimer, Emebet Adem, Endalkachew Nibret, Gizachew Yismaw, Edward Cruz Cervera, Kefale Ejigu, Dessalegn Tamiru, Abaineh Munshea, Ingrid Müller, Richard Weller, James A. Cotton, Pascale Kropf

## Abstract

Cutaneous leishmaniasis (CL) is a neglected tropical skin disease, caused by the protozoan parasite *Leishmania (L.)*. It is endemic to 90 countries and causes >200,000 new infections each year. In Ethiopia, CL is mainly caused by *L. aethiopica* and can present in different clinical forms: localised cutaneous leishmaniasis (LCL); mucocutaneous leishmaniasis (MCL), where the mucosa of the nose and/or the mouth are affected; and diffuse CL (DCL), characterised by non-ulcerating nodules. Persistent forms of LCL, as well as MCL and DCL, require treatment but are difficult to treat successfully and can lead to permanent disfigurement, social stigmatisation, and a mental health burden. The mechanisms behind the development of these different presentations of CL are not clearly understood, and both parasite and host factors could be involved. Here we analysed the whole genome sequence data for 48 clinical parasite isolates and show that parasites from CL cases with different presentations in a single Ethiopian setting are from the same genetic population. Furthermore, we did not identify any individual genetic variants significantly associated with disease presentation. We also measured plasma chemokine and cytokine levels of 129 CL patients presenting with different forms of CL. None of the cytokine or chemokine levels measured were significantly different between the different clinical presentations of CL. We also compared those with healthy nonendemic controls: our results show a chemokine but not a cytokine immune signature in patients with CL as compared to healthy nonendemic controls.

## INTRODUCTION

Cutaneous leishmaniasis (CL) is a neglected tropical disease caused by over 20 different species of the protozoan parasite *Leishmania (L.)* (1, 2). It is transmitted to mammalian hosts during the blood meal of infected sandflies. In Ethiopia, CL is mainly caused by *L. aethiopica* (3) with *Phebotomus (P.) longipes* and *P. pedifer* being the most common vectors (4, 5). The disease presents in three main clinical forms: Localised CL (LCL), where small nodules evolve into ulcerative lesions that usually heal within a few months; mucocutaneous leishmaniasis (MCL), where the lesions affect the nasal or oral mucosa; and diffuse CL (DCL), characterised by numerous non-ulcerating lesions. Both MCL and DCL, as well as persistent LCL rarely heal on their own and require treatment. CL often results in disfiguring scars and can lead to significant social stigma (6).

The mechanisms behind the development of these different presentations of CL are not well understood, and both parasite and host factors could be involved. There has been considerable interest in understanding how variation between parasites might result in differences in the outcome of *Leishmania* infections. The most obvious parasite genetic difference is that only a few *Leishmania* species are associated with visceral disease and a number of factors potentially involved in visceralisation have been identified (7). However, there are also clear differences between intraspecific parasite isolates and even between clones of a single isolate (8) in both *in vitro* growth and in infections of animal models (reviewed in (9)). In CL, peroxidase activity has been implicated in the dissemination of MCL-causing *L*. (*V*.) *guyanensis* strain in a rodent model (10). While most data suggests that parasite isolates from cutaneous and mucosal CL from the same patients are genetically very similar (11, 12), most of this work has relied on experimental characterisation of small numbers of parasite isolates taken from cases with differing clinical presentations. While whole-genome data from natural populations of many *Leishmania* species is now available (e.g (13–16)), there has been little work directly attempting to directly relate parasite genetic variation to clinical variation. One exception is a recent genome-wide association study that found some evidence that clinical outcome is linked to parasite genotype in *L. infantum* (16). Research on *L. aethiopica* is particularly neglected, and the only genomic data available for this species is from 19 historical isolates and some hybrid forms collected from cryopreserved culture collections (17). For these kinds of samples accurate epidemiological or clinical data is typically not available, and there is often significant variation in sampling times and places which is likely to make detecting clinical associations much more challenging. In addition to parasite factors, host factors, such as the skin microbiome (18); vector derived products (19); the genetics of the host (20); and immune responses (21) can influence CL development. In contrast to the experimental mouse model of CL (22–24), there are no clearly polarised Th1 and Th2 responses in patients with different clinical manifestations of CL, but a mixed production of cytokines: antigen-specific stimulation of peripheral blood mononuclear cells (PBMCs) from CL patients resulted in production of both Interferon (IFN)γ and Interleukin (IL)-4 during the active phase of the disease, but mainly IFNγ and low IL-4 after healing (25). There were also increased levels of Th1 type cytokines, such as IFNγ, IL-2 and Tumor Necrosis Factor (TNF)α; and Th2 type cytokines, such as IL-4, IL-5 and IL-13; as well as the regulatory cytokine IL-10 in the plasma of *L. guyanensis*-infected CL patients (26). In patients with mucosal leishmaniasis (ML), the antigen-specific production of IFNγ and IL-5 was higher as compared to patients with CL; and the detection of IL-4 was low and only present in some ML patients (27). In a study by Bacellar *et al.,* PBMCs from patients with ML produced increased levels of IFNγ and TNFα and decreased levels of IL-10 as compared to patients with CL (28). It is generally accepted that DCL patients have an inability to mount an efficient immune response (29). There is very limited information on the immune response in *L. aethiopica*-infected patients or on the mechanisms resulting in the development of different manifestations of CL. It has been shown that *L. aethiopica* parasites isolated from DCL and LCL patients induced different cytokine profiles in PBMCs (30). A recent study showed that in response to *L. aethiopica*, monocytes and neutrophils from LCL patients were more activated as compared to controls (31), however it was not compared between different clinical forms of CL.

The aim of this study was to assess whether different forms of CL were driven by genetic differences between the infecting parasites and whether they were associated with different host systemic immune signatures. To this end we recruited a cohort of CL patients in Nefas Mewcha Hospital, in Lay Gayint District, Northwest Ethiopia (32) and generated whole-genome sequence data from parasites isolated from these patients, as well as measured their plasma cytokine and chemokine profiles. This represents the most detailed immunological and clinical investigation of a large *L. aethiopica* CL cohort to date, and the first attempt to integrate an understanding of the parasite population with clinical and epidemiological data from the same cohort for this species.

## MATERIALS AND METHODS

### Ethical approvals and patients’ recruitment

This study was approved by the Research and Ethical Review Committee of the College of Science (RECCS), Bahir Dar University (RCSVD 002/2011), the National Research Ethics Review Committee (ref. No MoSHE/RD/ 14.1/10112/2020) of the Ministry of Science and Higher Education of Ethiopia and Imperial College Research Ethics Committee (ICREC 18IC4593). Informed written consent was obtained from each participant.

For this study, we analysed whole genome sequences of parasite isolates and plasma cytokines and chemokines from CL patients identified from the cohort that was described in (32). Patient recruitment, diagnosis and characteristics are described in (32). A further 22 healthy non-endemic controls (HNEC) were also recruited.

### Parasite isolates

A skin scraping was obtained from the edge of the lesion with a sterile scalpel and was added to culture medium (M199 medium with 25mM hepes, 0.2μM folic acid, 1mM hemin, 1mM adenine, 800μM Biopterin, 50 IU/mL penicillin, 50 mg/mL streptomycin and 10% fetal bovine serum (Sigma, USA)) and incubated at 26°C. Parasite growth was examined by microscopy. The promastigotes were then washed with PBS and the DNA was purified using the DNeasy Blood & Tissue Kits (Qiagen) following the manufacturers procedure. The DNA was stored at -20°C until further use.

### Genome sequencing and analysis of sequence data

Genomic DNA was isolated from parasite cell pellets using the QIAgen Blood and Tissue DNA kit. The isolated DNA was sheared into 400- to 600-bp fragments by focused ultrasonication (Covaris Inc.). Sequencing libraries were generated using a PCR-free approach (33) and then cleaned up using Agencourt AMPure XP SPRI beads (Beckman Coulter). The resulting libraries were sequenced on the Illumina NovaSeq platform at the Wellcome Sanger Institute. Low-quality bases at the 3’ end of reads and Illumina sequencing adaptors were removed using Trimmomatic v0.39 (34) with parameters “ILLUMINACLIP:PE_adaptors.fa:2:30:10 TRAILING:15 SLIDINGWINDOW:4:15 MINLEN:50”. Trimmed reads were then mapped against the reference genome of *L. aethiopica* L147 (35) obtained from TriTrypDB release 51 (36) using the mem mapper in the Burrows-Wheeler Aligner (bwa) version 0.7.17 (37). Reads were sorted and duplicates removed using samtools v1.14 and picard v 2.22.2 (https://broadinstitute.github.io/picard/). Nucleotide changes and small insertion-deletion variation from the reference were then identified using a pipeline based on the Broad Institute’s Genome Analysis ToolKit (GATK), generating per-sample gvcf files with *HaplotypeCaller* assuming diploid genotypes before jointly genotyping across the whole cohort. SNPs were then filtered using hard cut-offs for quality statistics with GATK “VariantFiltration” using the following filters: QD < 2.0; MQ < 50.0; FS > 20.0; SOR > 2.5; BaseQRankSum < -3.1; ClippingRankSum < -3.1; MQRankSum < -3.1; ReadPosRankSum < -3.1; and DP < 4. Coverage was calculated using samtools v1.17 (38) *depth* command and summaries per sample and per chromosome calculated using GNU datamash v1.2 and R v4.4.1. Samples were retained for further analysis if the median depth of coverage of at least 5x across the genome and sites were removed if more than 10% of the remaining samples had missing genotype calls for that sites.

Aneuploidy profiles for each sample were estimated by calculating the mean coverage across the genome, excluding chromosome 31, which was assumed to represent the diploid coverage. Per-chromosome mean coverage was divided by this diploid coverage and multiplied by 2 to estimate chromosome-specific coverage, and these estimates confirmed by inspection of the non-reference allele-frequency for each isolate and chromosome, inferred from the per-allele read depth estimates from GATK. Phylogenetic trees were reconstructed using whole-genome nucleotide variation between *L. aethiopica* samples by generating a diploid (heterozygous) consensus genome sequence for each sample by projecting SNP variants onto the reference genome using the *consensus* command in bcftools v1.14 (38), phylogenies were inferred using RAxML-NG v0.8.1(39) using the Jukes-Cantor substitution model. Trees were visualized using ggtree v3.8.2 (40) in R v4.4.3.

Population genomic analysis was based on SNP genotype calls for each sample. Principal-components analysis and tests for differences in genetic similarity within and between phenotype (clinical presentation) groups were performed in plink v1.90 (41). Tests for association between individual genetic variants and clinical presentation phenotypes were performed using *–assoc* (for chi-squared test of association) and *–logistic* (for logistic regression tests including PCA axes as explanatory variables) in plink v1.90. All other statistical analysis and plotting were performed in R v4.3.1, using tidyverse packages (42).

### Blood sample collection

Two ml of venous blood were drawn in heparin tubes and centrifuged, the plasma was collected and immediately frozen to be used at a later time point to measure the levels of cytokines and chemokines.

### Cytokine and chemokine measurements

IFNγ, IL-1β, IL-2, IL-4, IL-6, IL-8, IL-10, IL-12p70, IL-13, TNFα, Eotaxin, Eotaxin-3, Interferonγ-induced protein (IP)-10, monocyte chemoattractant protein (MCP)-1, MCP-4, macrophage-derived chemokines (MDC), macrophage inflammatory protein (MIP)-1α, MIP-1β and thymus- and activation-regulated chemokine (TARC) plasma levels were measured by multiplex assay using Cytokine and Chemokine panel V-PLEX Kits (Meso Scale Diagnostics, USA).

### Statistical analysis

Data were evaluated for statistical differences as specified in the legend of each table and figure. The following tests were used: Mann-Whitney, Kruskal-Wallis and Spearman. Differences were considered statistically significant at *p*<0.05. ∗=p<0.05, ∗∗=p<0.01, ∗∗∗=p<0.001 and ∗∗∗∗=p<0.0001. Unless otherwise stated, summary statistics given are medians followed by interquartile range (IQR) in square brackets.

## RESULTS

### Parasite genetic variation

We obtained high-quality whole genome sequence data for 48 isolates obtained from the cohort (Table S1). This *L. aethiopica* population showed a moderate level of genetic variation (196,277 variable SNP sites), and phylogenetic and principal components analysis of these data confirm that these parasites are closely related and form a monophyletic group distinct from other *Leishmania* species (figures 1B-D). As well as observing limited SNP variation between parasites in this population, we also see remarkably little variation in somy within the *L. aethiopica* population (Figure 2), with just occasional and sporadic trisomies, particularly for chromosome 1, and a few examples of chromosome loss. Chromosome 31 is observed to be generally tetrasomic, as is general in *Leishmania*. We see decay in linkage equilibrium over 10s of kilobases within this set of samples (Figure 1A) we find a high-to-moderate level of inbreeding in many isolates as frequently seen in *Leishmania* populations (full data not shown). Taken together, these results show that CL in Gayint is caused by a single, interbreeding population of *L. aethiopica*.

**Figure 1.**
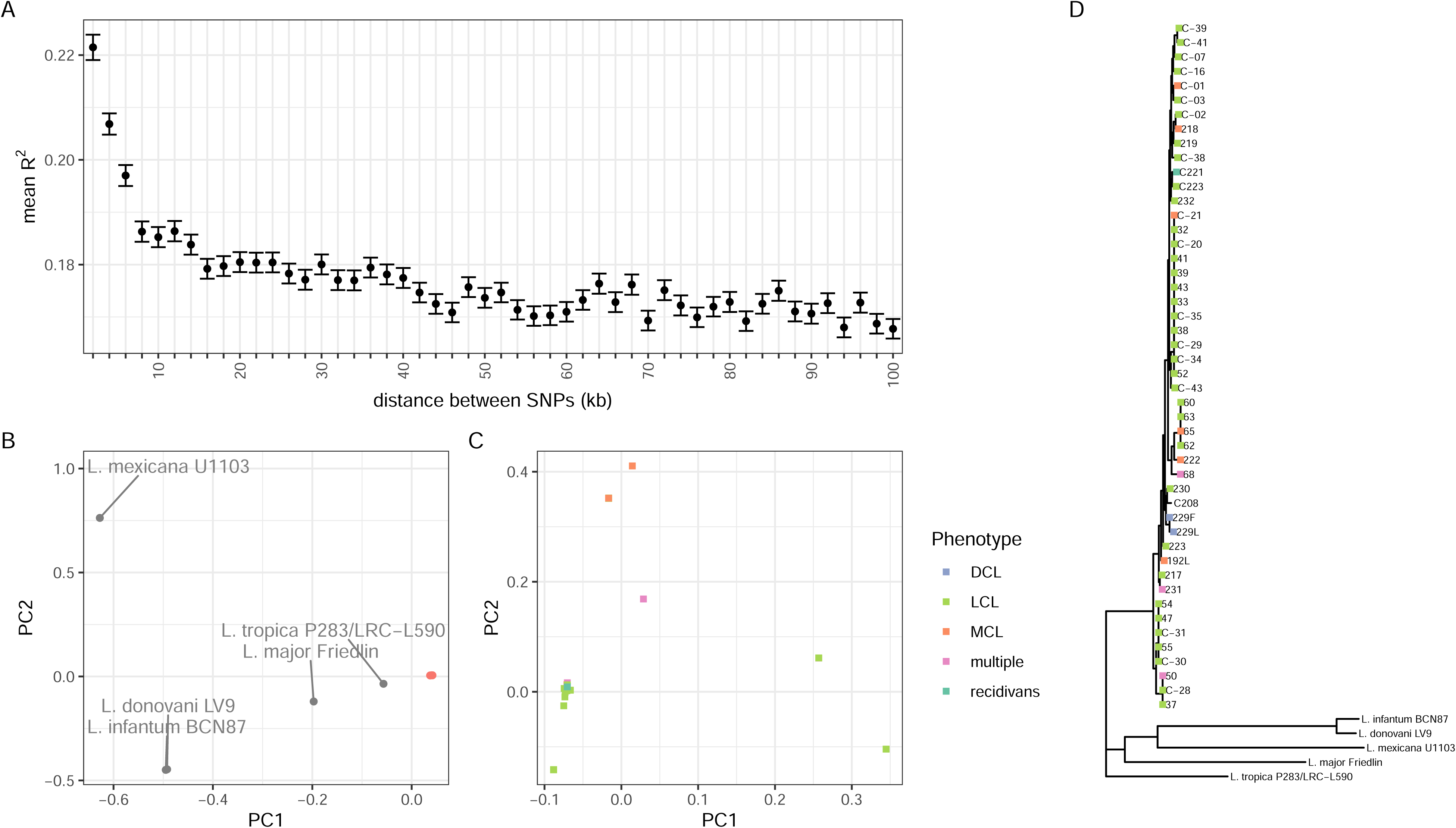
– Population genetics of Gayint isolates. (A) Decay of linkage disequilibrium with genomic distance. Symbols show mean R^2^ between pairs of 250,000 random SNPs on all chromosomes, and error bars show 1 standard error for variants in bins of 2kb distance centered on the indicated distance. R^2^ is calculated for 48 isolates from Gayint district. (B) and (C) Principal components analysis of isolates based on whole-genome SNP data of Gayint isolates (B) with and (C) without outgroups from 5 different *Leishmania* species. (D) Maximum-likelihood phylogeny of CL isolates from Amhara region and comparator isolates based on whole-genome SNP data. Coloured squares on leaves indicate clinical presentation phenotype. Scale bar is in expected number of substitutions per site.

**Figure 2.**
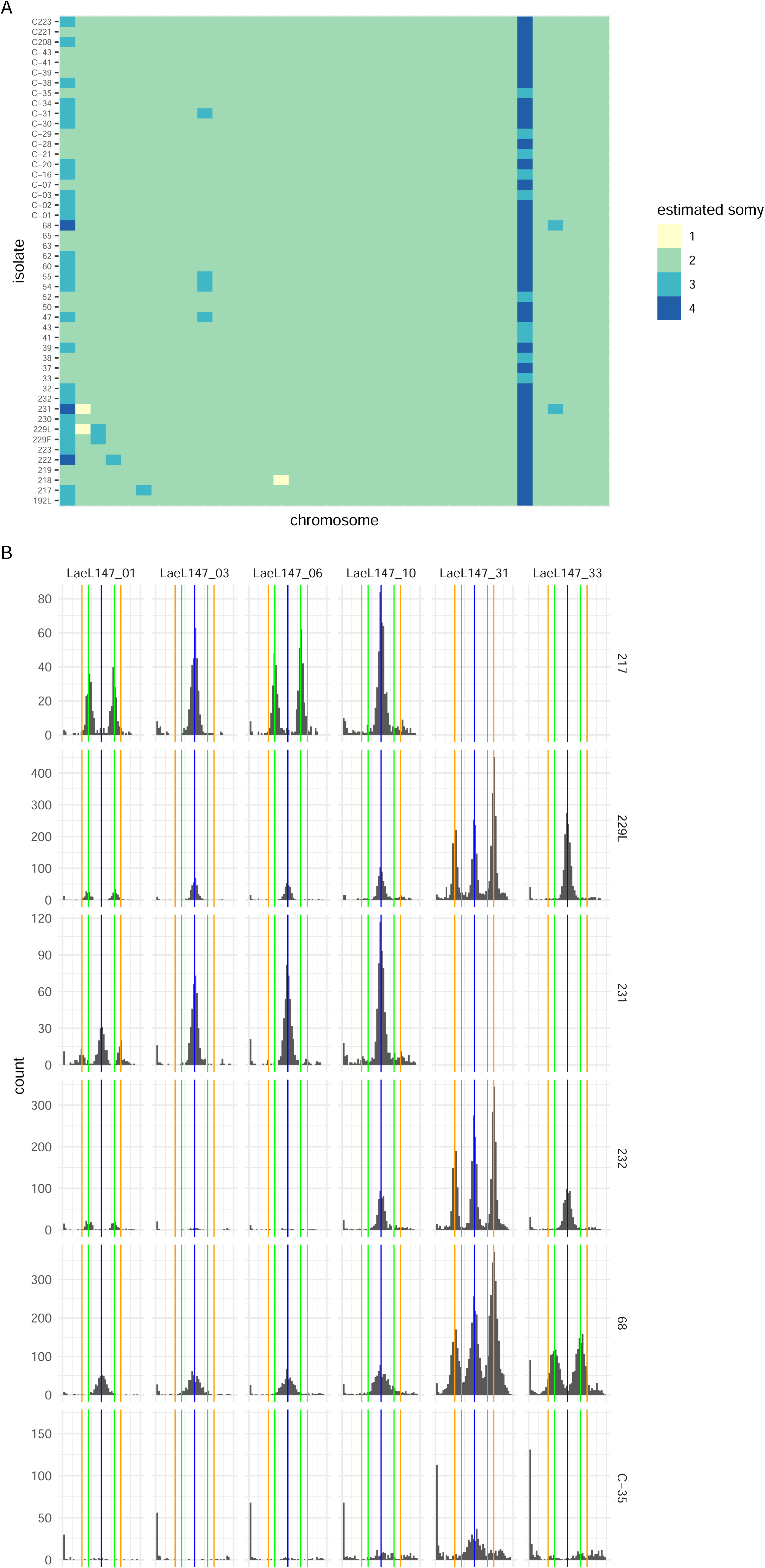
– Estimated somy. (A) Estimated chromosome copy number (somy) for all non-outgroup samples based on relative coverage and allele frequency. Rows represent individual isolates, columns each chromosome in order from 1 to 36 running left to right. Isolates are ordered by sampling location and clinical presentation. (B) example allele frequency distributions for selected chromosomes and isolates, supporting inferred trisomy and tetrasomies. Each panel is a histogram of non-reference allele frequencies for variants on a particular chromosome for one isolate. Blue vertical lines mark 0.5 allele frequency expected from disomy, green lines mark 1/3 and 2/3 allele frequencies expected from trisomy and orange lines mark ¼ and ¾ allele frequencies expected for tetrasomies.

### Genetic association with clinical presentation

Clinical phenotypes were available to classify 43 out of the 48 patients from which whole-genome data from parasite isolates was available as presenting with localised (35), mucosal (6) or disseminated (2) CL (4 other patients presented with multiple lesion types, and there was one case of leishmaniasis recidivans). The two parasites isolated from DCL patients are from lesions on the face and leg of the same patient and genetically very similar (99.97% identical SNP calls at polymorphic sites) – and so were considered likely to be from a single infection and no comparisons with DCL were attempted. There was no significant difference in genome-wide similarity between parasites isolated from patients with similar presentations to those isolated from patients with different presentations (Figure 3, Table 1). This confirms that the same genotypes of parasites are responsible for causing both disease presentations, rather than particular lineages being associated with specific presentations. For the 35 LCL isolates, we also classified 34 of these into contained (22) and spreading (12) LCL presentation (as described in (32) and again tested for whether parasites isolated from patients with particular forms of LCL were genetically distinct from parasites isolated from patients with different presentations. As before, no difference in overall genetic similarity within and between contained and spreading LCL was identified.

**Figure 3.**
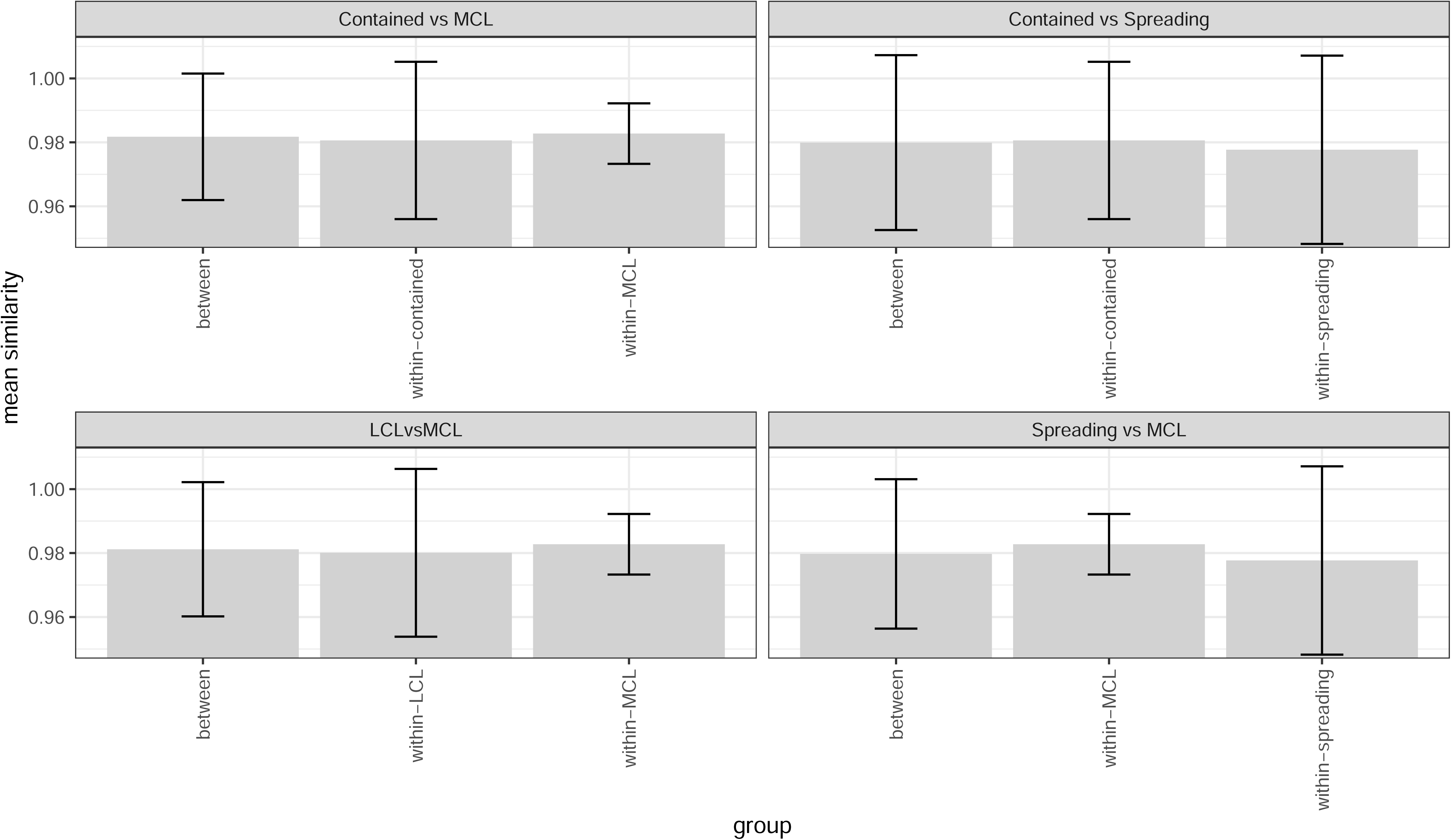
– Identity-by-similarity between Gayint isolates within and between groups of isolates with different presentations, based on whole-genome SNP variation data. Shaded bars represent mean similarity of all pairwise comparisons in each category, error bars represent 2 standard deviations. Note that y-axis scale does not start at zero.

**Table 1.** – Comparisons of genome-wide genetic similarity between parasites isolated from patients presenting with different forms of CL.

| **group / comparison** | **Identity-by-similarity** | **Standard deviation of IBS** | **p-value (test for greater similarity within group than between groups)** |
| --- | --- | --- | --- |
| LCL | 0.980065 | 0.0131195 | 0.627744 vs LCL-MCL |
|  |  |  | 0.50929 vs LCL-DCL |
| MCL | 0.98274 | 0.00473581 | 0.545355 vs LCL-MCL |
|  |  |  | 0.785182 vs MCL-DCL |
|  |  |  | 0.38874 (S LCL vs MCL) |
|  |  |  | 0.553924 (C LCL vs MCL) |
| LCL vs MCL | 0.98117 | 0.0104934 | N/A |
| S LCL | 0.977682 | 0.0147224 | 0.918641 (S LCL vs C LCL) |
|  |  |  | 0.689463 (S LCL vs MCL) |
| S LCL vs C LCL | 0.979914 | 0.0136714 | N/A |
| S LCL vs MCL | 0.979734 | 0.011677 | N/A |
| C LCL | 0.980597 | 0.0122999 | 0.312487 (S LCL vs C LCL) |
|  |  |  | 0.612314 (C LCL vs MCL) |
|  |  |  | 0.502915 (C LCL vs DCL) |
| C LCL vs MCL | 0.981731 | 0.00988552 | N/A |

Taking advantage of our genome-wide variation data, we also investigated whether individual single nucleotide variants were associated with particular presentations using a genome-wide association study (GWAS) design. For the comparison of LCL with MCL, a single significant SNP was identified at position 676,310 of chromosome 32 (figure S1A, Table 2). This site is intergenic, lying 73bp upstream of a putative DNA polymerase subunit (LAEL147_000639500). However, there was evidence that p-values from the GWAS were systematically inflated (figure S1B). This is likely to be due to some remaining population structure, or to linkage between SNP sites. Adjusting for these by including the first 2 principal component of genetic variation as covariates in the analysis removed the p-value inflation, but the identified SNP was no longer significant in this analysis. We have previously shown that LCL patients can be divided into two groups: those presenting with a well-defined contained lesion, with a distinct border around the lesion (contained LCL, C LCL) and those presenting with a lesion that did not have clear edges and was spreading (spreading LCL, S LCL) (32). We also compared parasites from the two LCL presentation types against MCL and against each other but found no significant associations with SNP variants in these comparisons.

**Table 2.** – Summary of genome-wide association results.

|  | Chi-squared test |  |  | Logistic regression (2 PCs) |  |  |
| --- | --- | --- | --- | --- | --- | --- |
| Comparison | Raw p-value of most significant variant | Corrected p-value of most significant variant | Number of SNPs with corrected p-value below 0.01 | Raw p-value of most significant variant | Corrected p-value of most significant variant | Number of SNPs with corrected p-value below 0.01 |
| LCL vs MCL | 2.79x10<sup>-9</sup> | 0.00052 | 1 | 0.004947 | 1 | 0 |
| S LCL vs C LCL | 0.00019 | 1 | 0 | 0.006731 | 1 | 0 |
| S LCL vs MCL | 1.66x10<sup>-5</sup> | 1 | 0 | 0.01816 | 1 | 0 |
| C LCL vs MCL | 1.66x10<sup>-5</sup> | 1 | 0 | 0.01011 | 1 | 0 |

We also tested for differences in chromosome copy number between parasites of different presentations, using a Fisher’s exact test to test for association between somy at each chromosome where there was variation in copy number and presentation phenotypes for each pair of presentations compared in the SNP-based GWAS. We found no significant associations (full results not shown).

### Chemokine and cytokine profile

Chemokines and cytokines were measured in the plasma collected from CL patients, that were part of the cohort described in (32), and from healthy nonendemic controls. 41 patients were females and 88 were males. The median age of the female CL patients was 32 [20.3-49] and that of males was 31 [20.3-45] years (p=0.9636, data not shown). The cohort of CL patients consisted of 96 LCL and 33 MCL patients. Of the 96 LCL patients, 65 presented with contained lesions (C LCL) and 31 with spreading lesions (S LCL). There were no significant differences between the duration of illness, the BMI or parasite grading between C LCL, S LCL and MCL patients (Table 3). However, S LCL patients presented with significantly more lesions as compared to C LCL and MCL patients (Table 3) (32).

**Table 3:** Clinical characteristics of CL patients.

|  | **C LCL (n=65)** | **S LCL (n=31)** | **MCL (n=33)** | **p-value** |
| --- | --- | --- | --- | --- |
| **Duration of illness (months)** | 9 [6-12] | 12 [7-12] | 9 [5-12] | 0.2750 |
| **BMI** | 21.1 [19.4-22.8] | 21.4 [19.1-22.7] | 20.0 [19.1-21.7] | 0.5146 |
| **Number of lesions** | 1 [1-1] | 2 [1-4] | 1 [1-1] | **0.0002** |
| **Parasite grading** | 1 [1-2] | 1 [1-2] | 1 [1-2] | 0.7844 |

### Chemokines

We first measured the levels of chemokines in the plasma of CL patients presenting with different clinical forms and compared them to those of healthy non endemic controls (HNEC). Whereas plasma levels of Eotaxin (p=0.2269), Eotaxin-3 (p=0.9983) and MCP-1 (p=0.5721) were similar between CL patients and controls, there were significantly higher levels of IL-8 (p<0.0001), IP-10 (p<0.0001), MCP-4 (p<0.0001), MDC (p<0.0001), MIP-1α (p=0.0019) MIP-1β (p<0.0001) and TARC (p<0.0001) in the plasma of CL patients as compared to HNEC (Figure 4). There were higher levels of MCP-4 in the plasma of patients with MCL and C LCL, but not S LCL and higher levels of MIP-1α in the plasma of C LCL, but not S LCL or MCL patients as compared to HNEC. No significant differences were observed between C LCL, S LCL and MCL patients (Table 4, Figure 4).

**Table 4:**
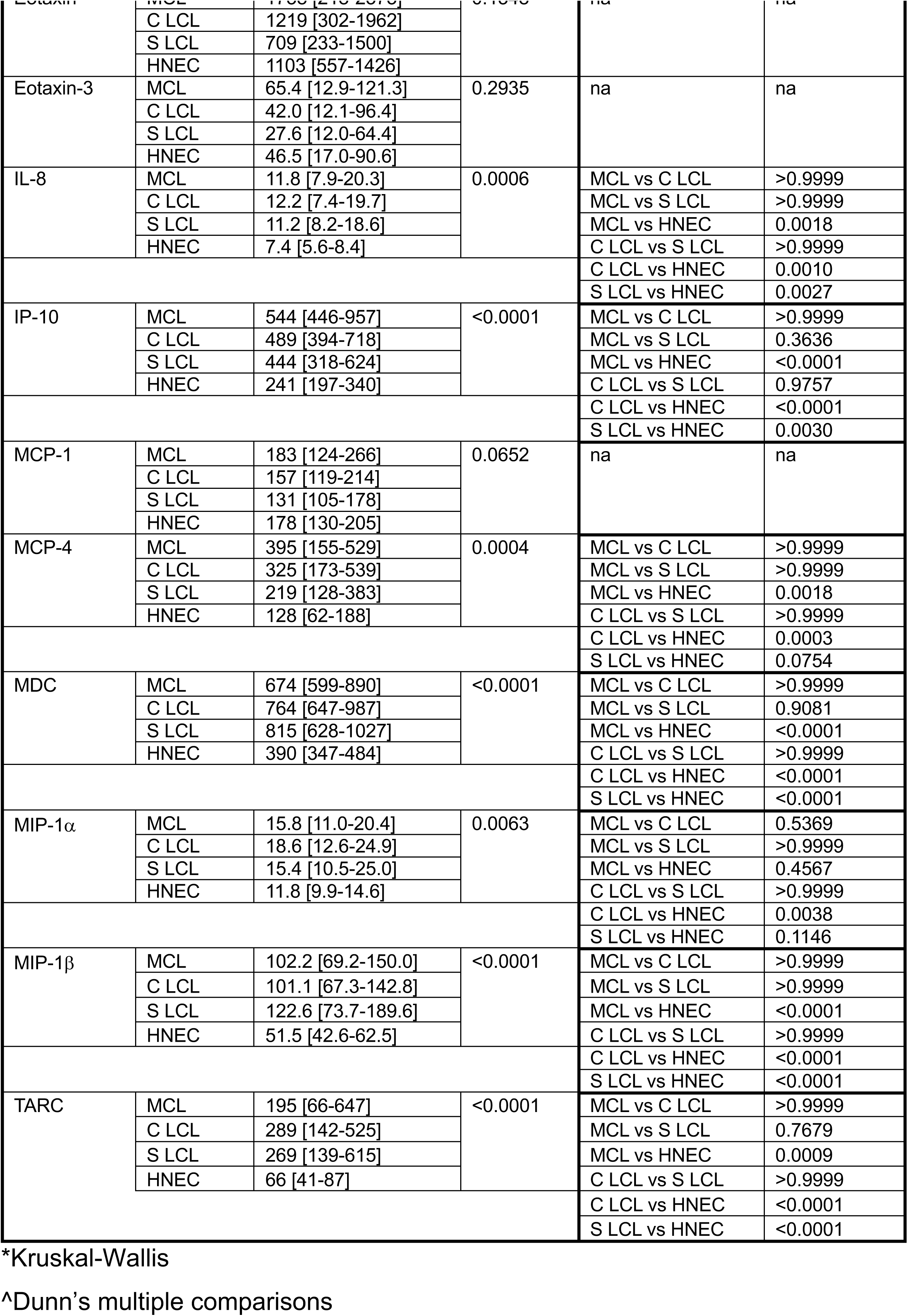
Comparison of plasma chemokine levels between CL patients presenting with different clinical forms and controls.

**Figure 4.**
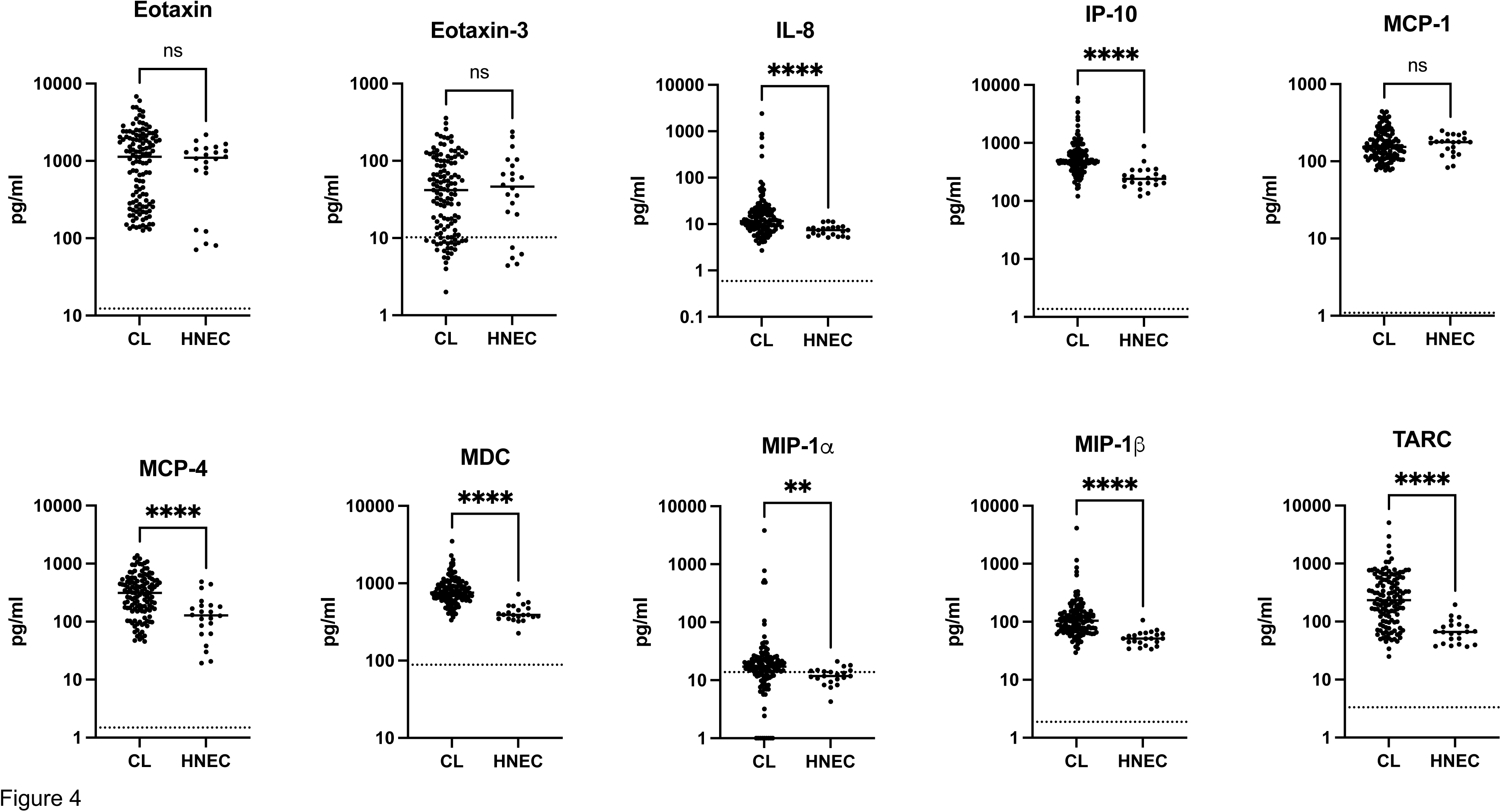
– Chemokine levels. Chemokine levels were measured in the plasma isolated from the blood of CL (n=127) and HNEC (n=22) by multiplex assay, as described in Materials and Methods. Each symbol represents the value for one individual, the straight lines represent the median. Statistical differences were determined using a Mann-Whitney test. CL= cutaneous leishmaniasis patients, HNEC=healthy non-endemic controls, ns=not significant.

### Cytokines

Of all the cytokines tested, only IFNγ and TNFα were above the detection limit (Figure 5). No significant differences were observed in plasma levels of IFNγ and TNFα between the different forms of CL patients and HNEC (Figure 5, Table 5). However, the levels of TNFα were significantly higher in LCL patients as compared to MCL patients (p=0.0359, Figure 5C).

**Figure 5.**
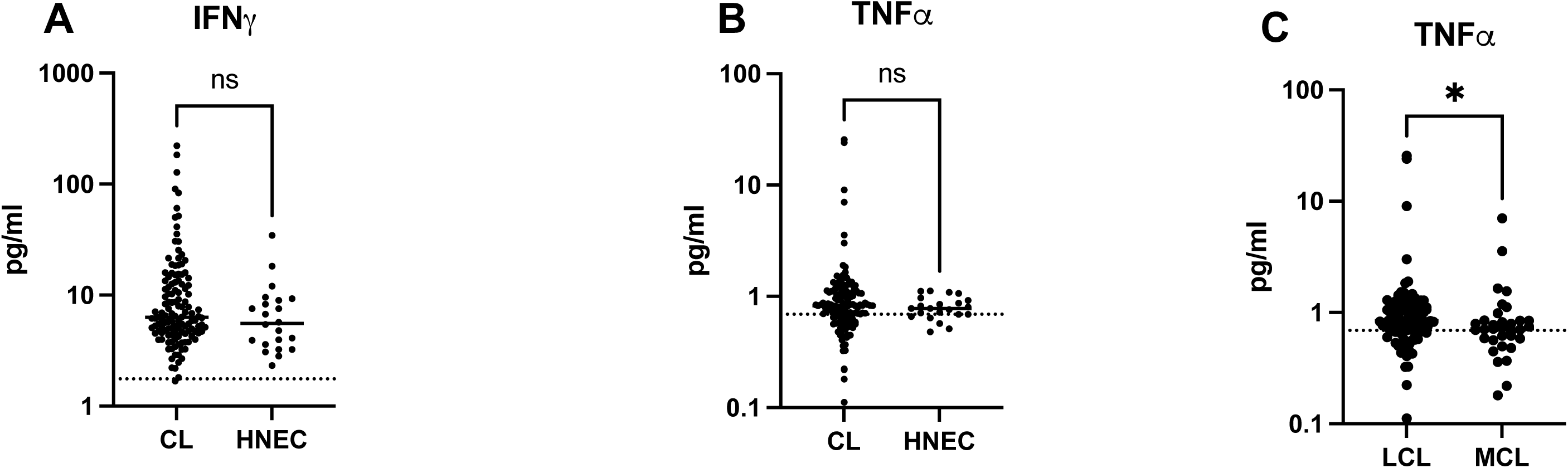
– Cytokine levels. Cytokine levels were measured in the plasma isolated from the blood of CL (n=129) and HNEC (n=22) by multiplex assay, as described in Materials and Methods. Each symbol represents the value for one individual, the straight lines represent the median. Statistical differences were determined using a Mann-Whitney test. CL= cutaneous leishmaniasis patients, HNEC=healthy non-endemic controls, ns=not significant.

**Table 5:** Comparison of plasma cytokine levels between CL patients presenting with different clinical forms and controls

|  |  |  | *p-value |
| --- | --- | --- | --- |
| IFNγ | MCL | 6.2 [3.8-10.3] | 0.2551 |
|  | C LCL | 7.0 [4.9-13.5] |  |
|  | S LCL | 6.2 [4.6-12.5] |  |
|  | HNEC | 5.6 [3.5-9.1] |  |
| TNFα | MCL | 0.7 [0.6-0.9] | 0.0903 |
|  | C LCL | 0.9 [0.7-1.2] |  |
|  | S LCL | 0.8 [0.7-1.1] |  |
|  | HNEC | 0.8 [0.7-0.9] |  |

Cytokine levels were measured in the plasma isolated from the blood of MCL (n=33), C LCL (n=65), S LCL (n=31) and HNEC (n=22) by multiplex assay, as described in Materials and Methods.
Each symbol represents the value for one individual, the straight lines represent the median. Statistical differences were determined using a Kruskal-Wallis test.
MCL=mucocutaneous, C LCL=contained localised cutaneous leishmaniasis, S LCL=spreading localised cutaneous leishmaniasis patients, HNEC=healthy non-endemic controls.

### Correlation between plasma chemokine and cytokine levels and clinical parameters

The clinical parameters collected from these patients were wide-ranging: the age of the CL patients varied from 18 - 68 years; the BMI from 16.3 - 27.3 kg/m^2^; the numbers of lesions from 1 - >5; the parasite gradings from 1+ - 6+; and the duration of illness from 1 -180 months. Therefore, we investigated if there was any association between these parameters and the plasma cytokines and chemokines. Correlations were observed between age and the levels of IP-10 and MCP-1 in C LCL patients; and between the parasite grading and Eotaxin, Eotaxin-3 and MCP-1 in MCL patients (Table 6). None of the other correlations were significant (data not shown).

**Table 6:** correlation between chemokine levels and different clinical parameters

|  |  | p-value | r |
| --- | --- | --- | --- |
| Age/IP-10 | C LCL | 0.0089 | 0.3290 |
| Age/MCP-1 | C LCL | 0.0125 | 0.3129 |
| Parasite grading/Eotaxin | MCL | 0.0120 | -0.5041 |
| Parasite grading/Eotaxin-3 | MCL | 0.0040 | -0.5650 |
| Parasite grading/MCP-1 | MCL | 0.0112 | -0.5081 |

Correlations between age and IP-10 and MCP1 (n=65), and parasite gradings and Eotaxin, Eotaxin 3 and MCP-1 (n=33) were measured by Spearman test.

## DISCUSSION

Here, we present the first whole-genome data from recent clinical isolates of *L. aethiopica* sampled in a well-described epidemiological context and accompanied by clinical and immunological data. Across *Leishmania* species, genomic data is only available for a few collections of recent clinical isolates (e.g. 14, 16, 43, 44). The only previously available genomic data for *L. aethiopica* apart from the reference genome assembly (35) was from historical cryopreserved isolates (17), although microsatellite and RFLP data from small numbers of isolates has been published (45, 46). We find limited genetic diversity across the set of isolates, although this is still very much more diverse than the largely clonal population responsible for most VL cases in the Indian subcontinent (44) but less so than in the *L. braziliensis* species complex (14). We find rapid decay in linkage disequilibrium within these isolates (as found across the species (17)). These isolates thus likely represent a single, interbreeding population of related *L. aethiopica* strains causing cutaneous leishmaniasis in Gayint: we find no evidence that other *Leishmania* species are involved in CL in Gayint as has been reported in Ethiopia (47) and elsewhere in Africa (48); Kenya (49) and in Yemen (50).

As in previous work (46), we find no significant genetic differentiation between parasites causing different clinical presentations of CL. The availability of genome-wide variation data allowed us to test for association between single nucleotide variants and clinical presentations and we also did not identify a convincing signature of association between any single nucleotide variation and clinical presentation in a genome-wide association test with these data. The single intergenic SNP significantly associated with MCL in contrast to LCL cases did not remain significant when attempting to correct for population structure within this set of isolates. In any case, the functional relevance of a non-coding SNP variant is unclear, even if this is sufficiently close to the downstream gene (a putative DNA polymerase subunit) to be part of the 5’ UTR and thus have a potential role in mediating gene expression (51); although the role of these elements is much less well-defined than for 3’ UTRs. While we might expect a more profound parasite difference between the relatively localised infections in LCL and MCL and the more widely disseminating DCL, the low prevalence of DCL in this setting (32) makes it challenging to assemble a sufficiently large patient cohort to identify variation associated with this presentation.

Our results identify a chemokine signature in patients with CL as compared to HNEC: we show that the levels of IL-8, IP-10, MCP-4, MDC, MIP-1α, MIP-1β and TARC are higher in the plasma of CL patients as compared to controls. This is in agreement with the study by de Mesquita *et al.* that showed that in patients with *L. guyanensis* infection, Eotaxin, IL-8, IP-10, MCP-1, MIP-1α, MIP-1β were increased as compared to controls; and that of Vargas-Inchaustegui *et al.*, that showed that IP-10 and MIP-1β were also increased in patients infected with *L. braziliensis* (52). Our results also show that there were no differences in plasma chemokine levels between C LCL, S LCL and MCL patients. However, as compared to HNEC, S LCL patients had similar levels of MCP-4 and S LCL and MCL patients had similar levels of MIP-1α; suggesting that there are differences in some chemokine levels between HNEC and CL patients with different clinical presentations. The increased chemokine levels we show in this study could be indicative of both wound healing or uncontrolled inflammation (53–55).

We also show that there were no differences in IFNγ and TNFα levels in the plasma of CL patients and controls. This is in contradiction to different studies, e.g. Castellano et al. (56), Espir et al. (57) and de Mesquita et al. (26) that showed high levels of these cytokines during active CL. These differences might be due to differences in infecting parasites, in the number of patients tested and importantly, to differences in clinical characteristics of the CL patients. In most cases, CL patients from studies such as (26, 52, 56, 57) present with a variety of different ages. Often, the CL cohorts are not characterised in detail, with no information about their BMI, duration of illness or number of lesions. The classification of the type of lesions is also complex. Our results show a lot of heterogeneity in patients age, in the duration of their illness, the number of lesions and their BMI. These are all factors that might influence the immune response. For example, TNFα (58), IFNγ (59), IP-10 (60) and MCP-1 (61) have been shown to increase with age. And indeed, we found positive correlation between age and MCP-1 and IP-10 in C LCL patients. The levels of cytokines and chemokines are also likely to be influenced by the duration of illness: a study by Melby *et al.* showed increased levels of IL-1α, TNFα, IL-10, and TGFβ mRNAs in older lesions (62). The nutritional status of the host also impacts on the levels of cytokines: we have previously shown that malnutrition results in altered cytokine profiles, with negative correlations between BMI and cytokines such as TNFα and IFNγ (63).

We also found negative correlations between the parasite gradings and Eotaxin, Eotaxin-3 and MCP-1 in MCL patients. MCL lesions are characterised by low parasite numbers (64) and indeed, the grading in the large majority of the lesions in our study were 1+ and 2+, with only one patient with a 3+ and one patient with a 4+. It also should be noted that the parasite grading might not be an accurate representation of the number of parasites in the lesions, as it is not been established whether parasites are distributed equally at the border of the lesions, where the scraping are collected.

Our study highlights that many parameters are likely to influence the plasma chemokine and cytokine levels in CL patients. To assess whether a systemic immune signature is associated with different clinical presentation of CL, large studies need to be undertaken with a precise characterisation of the CL cohorts and a longitudinal follow-up. Similarly, if parasite genetic factors are responsible for differences in presentation, larger cohorts – particularly of the rarer presentations – will be needed to detect these. Recruiting such cohorts in Ethiopia will be challenging, with CL affecting predominantly difficult to access rural communities in highland areas (65).

The differences between our results and those from other studies of CL might suggest a difference between *L. aethiopica* CL and disease caused by other *Leishmania* species, with most other research being on *L. braziliensis* or other neotropical *Leishmania* species from the subgenus *Viannia*. By establishing either immunological differences between healing and persistent lesions, or genetic markers for parasites likely to cause persistent lesions it might be possible to identify patients that are likely to develop severe persistent CL lesions early and so ensure that these patients receive early treatment.

## ACKNOWLEGEMENTS

The authors are thankful to staff of Nefas Mewcha Hospital for their enthusiastic collaboration during the data collection of this study.

This research is jointly funded by the UK Medical Research Council (MRC) and the Foreign Commonwealth and Development Office (FCDO) under the MRC/FCDO Concordat agreement (MR/R021600/1) (EY, BG, MY, SF, JAC, PK). JAC is funded by Wellcome via core funding of the Wellcome Sanger Institute (grant 206194).

---

**Figure S1.**
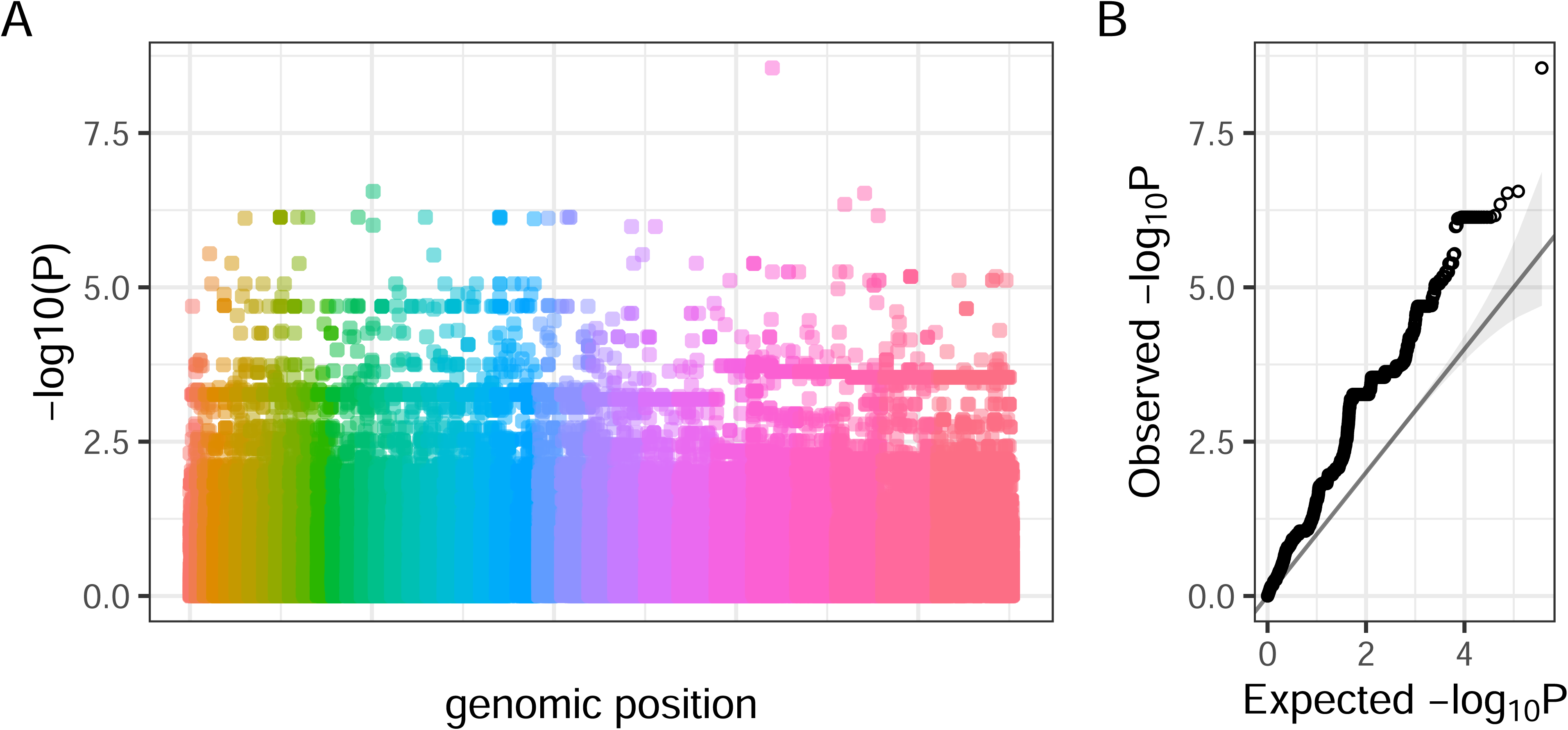
– Genome-wide association between LCL and MCL presentation phenotypes and SNP variants in Gayint *L. aethiopica*. (A) Manhattan plot of p-values for association between LCL vs MCL presentation and SNP variants. Each point represents a single SNP variation, with position on x-axis indicating position on genome. Colours distinguish SNPs on different chromosomes. (B) QQ-plot of -ve log (base 10) of expected p-value under null hypothesis vs observed p-values. Each point represents association between a SNP variant and LCL vs MCL presentation phenotype as shown in panel A, ordered by significance.

