## Supplemental table for "Parasite genetic variation and systemic immune responses are not associated with different clinical presentations of cutaneous leishmaniasis caused by *Leishmania aethiopica*"

**Table S1 – Samples used in genetic analysis**

| Sample name | Phenotype | Number of pairs of reads generated | Total Sequence Data generated (bp) | ENA sample accession number |
|---|---|---|---|---|
| 32 | LCL | 17516120 | 2644934120 | ERS6367640 |
| 33 | LCL | 15088118 | 2278305818 | ERS6367626 |
| 37 | LCL | 16390578 | 2474977278 | ERS6367642 |
| 38 | LCL | 17555242 | 2650841542 | ERS6367651 |
| 39 | LCL | 19556730 | 2953066230 | ERS6367649 |
| 41 | LCL | 15855062 | 2394114362 | ERS6367655 |
| 43 | LCL | 16809078 | 2538170778 | ERS6367641 |
| 47 | LCL | 14953826 | 2258027726 | ERS6367633 |
| 50 | multiple | 19614396 | 2961773796 | ERS6367643 |
| 52 | LCL | 14918932 | 2252758732 | ERS6367647 |
| 54 | LCL | 17937616 | 2708580016 | ERS6367632 |
| 55 | LCL | 17550922 | 2650189222 | ERS6367634 |
| 60 | LCL | 19896486 | 3004369386 | ERS6367639 |
| 62 | LCL | 18447182 | 2785524482 | ERS6367645 |
| 63 | LCL | 17841260 | 2694030260 | ERS6367638 |
| 65 | MCL | 17136990 | 2587685490 | ERS6367652 |
| 68 | multiple | 17960148 | 2711982348 | ERS6367648 |
| C-01 | MCL | 16710894 | 2523344994 | ERS6367631 |
| C-02 | LCL | 18167926 | 2743356826 | ERS6367658 |
| C-03 | LCL | 18733882 | 2828816182 | ERS6367636 |
| C-07 | LCL | 21415936 | 3233806336 | ERS6367653 |
| C-16 | LCL | 18708912 | 2825045712 | ERS6367657 |
| C-20 | LCL | 24597900 | 3714282900 | ERS6367629 |
| C-21 | MCL | 20091276 | 3033782676 | ERS6367650 |
| C-28 | LCL | 17845426 | 2694659326 | ERS6367627 |
| C-29 | LCL | 18579430 | 2805493930 | ERS6367635 |
| C-30 | LCL | 16151408 | 2438862608 | ERS6367628 |
| C-31 | LCL | 16362374 | 2470718474 | ERS6367644 |
| C-34 | LCL | 20844586 | 3147532486 | ERS6367646 |
| C-35 | LCL | 20128524 | 3039407124 | ERS6367630 |
| C-38 | LCL | 15115736 | 2282476136 | ERS6367637 |
| C-39 | LCL | 17201404 | 2597412004 | ERS6367625 |
| C-41 | LCL | 17521432 | 2645736232 | ERS6367656 |
| C-43 | LCL | 17283186 | 2609761086 | ERS6367654 |
| 232 | LCL | 48800472 | 7368871272 | ERS13501735 |
| 229L | DCL | 55626720 | 8399634720 | ERS13501736 |
| 218 | MCL | 46393864 | 7005473464 | ERS13501737 |
| 217 | LCL | 45617724 | 6888276324 | ERS13501738 |
| 192L | MCL | 52175738 | 7878536438 | ERS13501739 |
| C221 | recidivans | 45224654 | 6828922754 | ERS13501725 |
| 229F | DCL | 52126450 | 7871093950 | ERS13501726 |
| 230 | LCL | 50540918 | 7631678618 | ERS13501728 |
| 231 | multiple | 47430720 | 7162038720 | ERS13501729 |
| C208 | multiple | 47544092 | 7179157892 | ERS13501730 |
| 223 | LCL | 46383638 | 7003929338 | ERS13501733 |
| C223 | LCL | 44568610 | 6729860110 | ERS13501731 |
| 222 | MCL | 38334880 | 5788566880 | ERS13501732 |
| 219 | LCL | 45523200 | 6874003200 | ERS13501734 |
